# The regulation of Protein Phosphatase 4 by FBXO42 is required for cancer cell survival

**DOI:** 10.1101/2025.04.22.649889

**Authors:** Stephan H. Spangenberg, Nicholas Garaffo, Enya Marie Catherine Alcindor, Ben Lusk, Vijaya Pandey, Adam Longhurst, Emma Bolech, Becky Xu Hua Fu, Luke A. Gilbert, James A. Wohlschlegel, David Toczyski

## Abstract

FBXO42 is a poorly characterized F-box protein that is essential in 15% of cancer cell lines from diverse tissue types. FBXO42 has been implicated in the regulation of mitosis and p53 signaling. High-throughput approaches indicate that FBXO42 function correlates with that of CCDC6, and that the two proteins interact physically, but the relationship between these proteins is not understood. Through a genome-wide CRISPR knockout screen, we found that mutation of FBXO42 is synthetically lethal with mutations in the γ- tubulin ring complex proteins MZT1 and MZT2B, suggesting that cells with centrosome and/or mitotic spindle assembly dysfunction are more sensitive to FBXO42 loss. Furthermore, we found that FBXO42 and CCDC6 contribute to p53 activation in response to centrosome depletion. Using mass spectrometry-based proteomics, we found that FBXO42 binds, is required for the ubiquitination of, and negatively regulates the expression of PPP4C (protein phosphatase 4 catalytic subunit). FBXO42’s interaction with PPP4C was independent of CCDC6. Similarly, we found that CCDC6 physically interacts with PPP4C independently of FBXO42 and does not affect PPP4C ubiquitination. Knockdown of PPP4C reduced FBXO42-CCDC6 interactions, suggesting that FBXO42 and CCDC6 may bind to and regulate PPP4C through separate mechanisms. Using gene knockdown rescue experiments, we confirmed that aberrant expression of PPP4C is a major driver of cell death in an FBXO42-essential Neuroblastoma cell line. These findings shed light on the function of two poorly understood proteins in regulating PP4 activity, p53 signaling, mitosis and cancer cell survival. A better understanding of FBXO42 and CCDC6 could inform the development of targeted cancer therapeutics.

## INTRODUCTION

FBXO42 is a substrate receptor for the SCF, an E3 ubiquitin ligase composed of the Ring finger protein RBX1, the cullin CUL1, and an adaptor protein SKP1, which binds F-box proteins that serve as substrate receptors (Bai et al., 1996; Bennett et al., 2010; Craig and Tyers, 1999; Deshaies and Joazeiro, 2009; Frescas and Pagano, 2008; Grim et al., 2008; Jin et al., 2004; Kipreos and Pagano, 2000; Liwocha et al., 2024; Rape et al., 2006; Skaar et al., 2009; Skowyra et al., 1997; Tyers and Willems, 1999; Zheng et al., 2002). FBXO42 is essential in approximately 15% of cancer cell lines, including cancer types from diverse lineages, but is not essential in non-cancerous cells such as RPE1 (Hoellerbauer et al., 2024; Toledo et al., 2015; Tsherniak et al., 2017; Zhou et al., 2024). There is debate in the literature as to whether FBXO42 loss affects p53 levels, with one study seeing only an effect on the stability of particular mutant alleles (Lü et al., 2024). It was previously shown that FBXO42 is important for mitosis and correct assembly of the mitotic spindle, although the mechanism of this regulation is unknown (Hoellerbauer et al., 2024; Hundley et al., 2021). FBXO42 has also been implicated in acquired resistance to MEK1/2 inhibitors (Nagler et al., 2020); Notch signaling (Jiang et al., 2022); correct synaptonemal complex assembly and the unfolded protein response in Drosophila (Barbosa et al., 2020; Santos et al., 2024). Despite its implication in these important pathways, we still lack a comprehensive understanding of the function and substrates of FBXO42, or why it is essential in a subset of cancer cells.

The DepMap cancer dependency database (https://depmap.org/portal) shows that FBXO42 has a very high co-dependency correlation with the coiled-coil domain protein CCDC6 (Tsherniak et al., 2017). FBXO42 and CCDC6 have been reported to function together in regulating USP28 and p53 (Lü et al., 2024), and they physically interact based on high throughout proteomics analyses (Huttlin et al., 2021; Huttlin et al., 2017; Oughtred et al., 2021). CCDC6 plays a role in the DNA damage response (Morra et al., 2022), and commonly forms an oncogenic fusion protein with RET (Cerrato et al., 2018; Li et al., 2019). However, the function of CCDC6 is not well-understood.

In this study, we investigated the function of FBXO42 and CCDC6 in cancer cell survival and mitosis. Our data showing a genetic interaction between FBXO42 and the γ-tubulin ring complex are consistent with previous findings that FBXO42**^-/-^** cells have higher frequencies of monopolar spindles (Hundley et al., 2021). We find that FBXO42 depleted cells are sensitive to centrosome dysfunction, and that FBXO42 and CCDC6 promote p53 activation following centrosome depletion. We identified PPP4C, the catalytic subunit of protein phosphatase 4 (PP4), as a ubiquitination substrate of FBXO42. PP4 is involved in numerous aspects of cellular physiology including mitotic spindle assembly and γ- tubulin function, as well as the DNA damage repair (Park and Lee, 2020; Toyo-oka et al., 2008; Voss et al., 2013), and is localized to centrosomes (Martin-Granados et al., 2008; Voss et al., 2013). Knockdown of PPP4C partially rescued cell death/arrest caused by FBXO42 or CCDC6 knockdown in the glioblastoma cell line LN-18, suggesting that PPP4C is the critical target of these proteins. Finally, our data suggest that CCDC6 and FBXO42 downregulate PPP4C activity via distinct mechanisms.

## RESULTS

### CRISPR/Cas9 Screen for genes that confer sensitivity or resistance to FBXO42 knockout

To explore the function of FBXO42, we performed a CRISPR/Cas9, genome-wide knockout screen (Figure 1A) in WT or FBXO42^-/-^ HAP1 cells. Cells were untreated (Figure 1B), or treated with compounds that disrupt mitosis, to which FBXO42^-/-^ cells have heightened sensitivity (Hundley et al., 2021): BI-2536 (Figure 1C), a PLK1 inhibitor, or Colchicine (Figure 1D), an inhibitor of microtubule polymerization (Dalbeth et al., 2014). Amongst genes whose mutation sensitized cells to FBXO42 loss in all three conditions were MZT1 and MZT2B, members of the γ-tubulin ring complex (Figure 1E). Interestingly, FBXO42 knockout partially rescued the effects of epsilon-tubulin (TUBE1) knockout (Figure 1E). We verified these results by transducing WT or FBXO42^-/-^ HAP1 cells expressing Cas9 with lentivirus constructs encoding sgRNA against MZT1, MZT2B, or TUBE1. We observed that knockout of TUBE1 reduced cell proliferation/survival, as previously seen (Costanzo et al., 2016). FBXO42 knockout cells showed resistance to TUBE1 knockout most strongly in BI-2536-treated cells, whereas FBXO42 mutation was synthetically lethal with MZT1 or MZT2B knockout in both BI-2536 treated and untreated cells (Figure 1F and 1G). It has been observed that TUBE1 knockout causes impaired centriole stability and arrest of cell division in a p53-dependent manner. With this in mind, we investigated the role of FBXO42 in p53 signaling (Wang et al., 2017).

**Figure 1:**
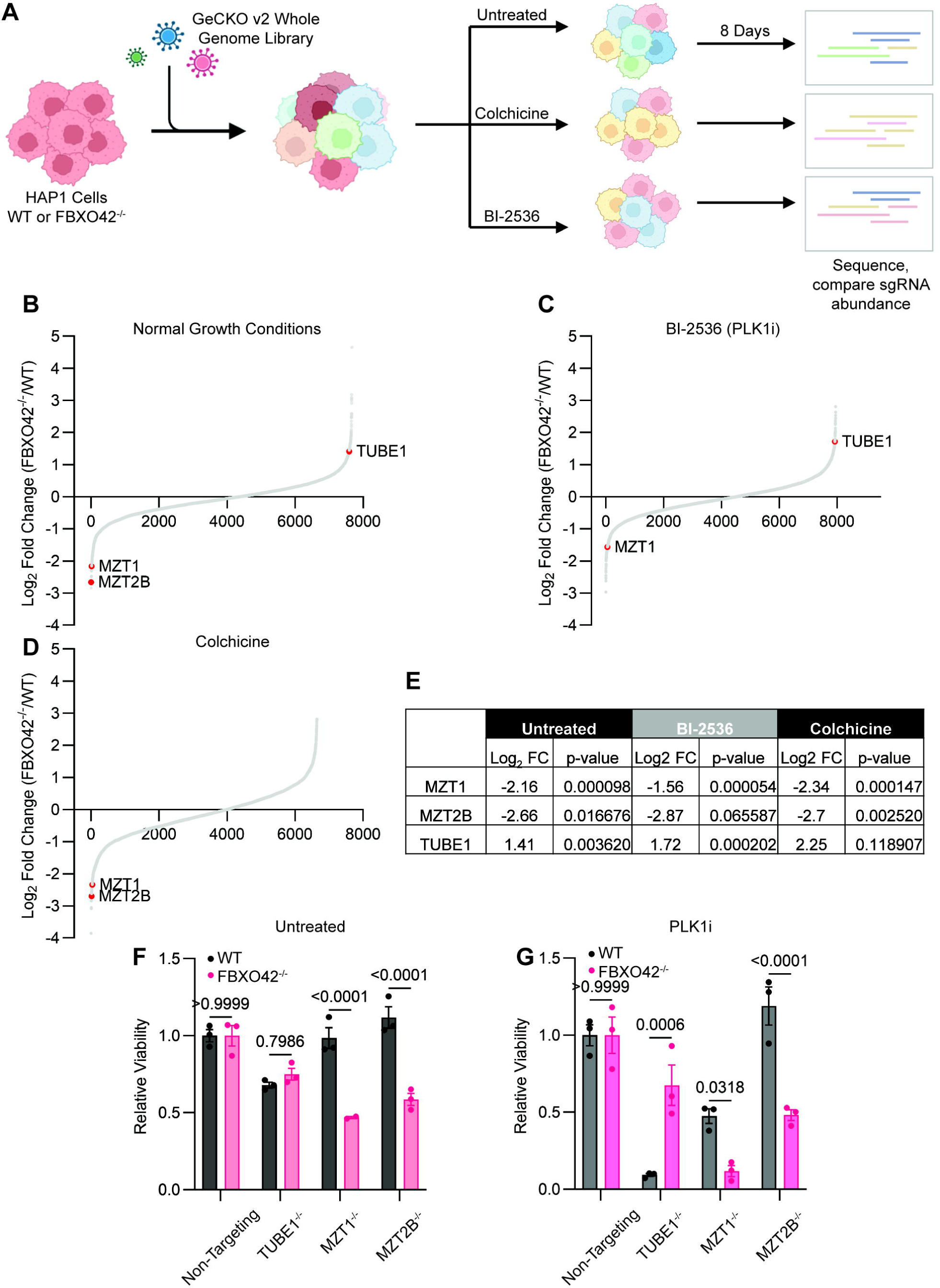
CRISPR/Cas9 Screen for genes that confer sensitivity or resistance to FBXO42 knockout. **A.** Schematic of a genome wide CRISPR knockout screen to identify genes that sensitized or conferred resistance to FBXO42 knockout cells. This screen was performed under normal growth conditions or with colchicine (10 nM) or BI-2536 (3 nM) for eight days in duplicate. **B-D.** Overview of CRISPR screen results. Log2 fold-change enrichment or de-enrichment of sgRNAs targeted genes are shown. Only sgRNA hits with an adjusted p-value less than 0.05 are included. Some genes which were common hits in multiple screens are highlighted. **E.** Log2 fold-change enrichment or de-enrichment and Mann-Whitney p-values for selected genes. FBXO42 knockout cells compared to WT cells. **F.** Crystal violet-based viability assay showing the relative viability of HAP1 cells, WT or clonal FBXO42 knockout. TUBE1, MZT1, or MZT2B were knocked out using induced expression of CRISPR-Cas9, and viability was assessed a total of 13 days later. Viability is normalized to the non-targeting control of either WT or FBXO42^-/-^ cells. Each point is a replicate well. Two-way ANOVA. p-value of individual comparisons, with multiple comparison correction, is displayed. **G.** As **F**, but with cells treated with 3 nM BI-2536, starting three days after knockout of MZT, MZT2B, or TUBE1.

### FBXO42 and CCDC6 are Involved in p53 Activation Following Centrosome Dysfunction

DepMap shows a positive correlation between FBXO42 and TP53, and a negative correlation between FBXO42 and negative regulators of p53 (Tsherniak et al., 2017) (https://depmap.org/portal) (Figure 2A-B); the same subset of cell lines that grow at increased rates when TP53 is deleted also grow at increased rates when FBXO42 is deleted. However, FBXO42 is essential in many cell lines, and this occurs at a higher frequency in TP53 mutant cells than in TP53 WT cells (Figure 2A). This suggests that FBXO42 knockout causes cell death or arrest, at least partially, through p53-independent mechanisms. To elucidate the role of FBXO42 in cells with an intact p53 pathway, we generated FBXO42 knockout clones of hTERT immortalized retinal pigment epithelial cells (RPE1) (Figure 2C, S2A). We also knocked out CCDC6 (Figure 2C, S2B), which has a very high co-dependency correlation with FBXO42 in regards to cell line essentiality (Figure 2B). Both FBXO42^-/-^ and CCDC6^-/-^ RPE1 cells had increased sensitivity to the PLK1 inhibitor BI-2536 compared to WT cells (Figure 2D), consistent with previous results for FBXO42 in HAP1 cells (Hundley et al., 2021). They did not differ in sensitivity to rotenone, which kills cells via an unrelated mechanism. Considering that FBXO42 mutation reduced cell death caused by loss of TUBE1, and that TUBE1 knockout has been shown to inhibit centriole duplication, we hypothesized that FBXO42 may be involved in p53 activation following centrosome depletion (Bazzi and Anderson, 2014; Fong et al., 2016; Lambrus et al., 2016; Meitinger et al., 2016; Wong et al., 2015). We treated cells with the PLK4 inhibitor centrinone B, which is known to activate p53 signaling via depletion of centrosomes, without induction of DNA damage (Denu et al., 2018; Pei et al., 2024; Rodríguez-Real et al., 2023; Singh et al., 2022; Wong et al., 2015). We found that FBXO42 and CCDC6 knockout cells had reduced induction of p53 and p21 following centrinone B treatment (Figure 2E, 2F). This suggests that FBXO42 and CCDC6 are required for the activation of p53 in response to inhibition of centrosome duplication.

**Figure 2:**
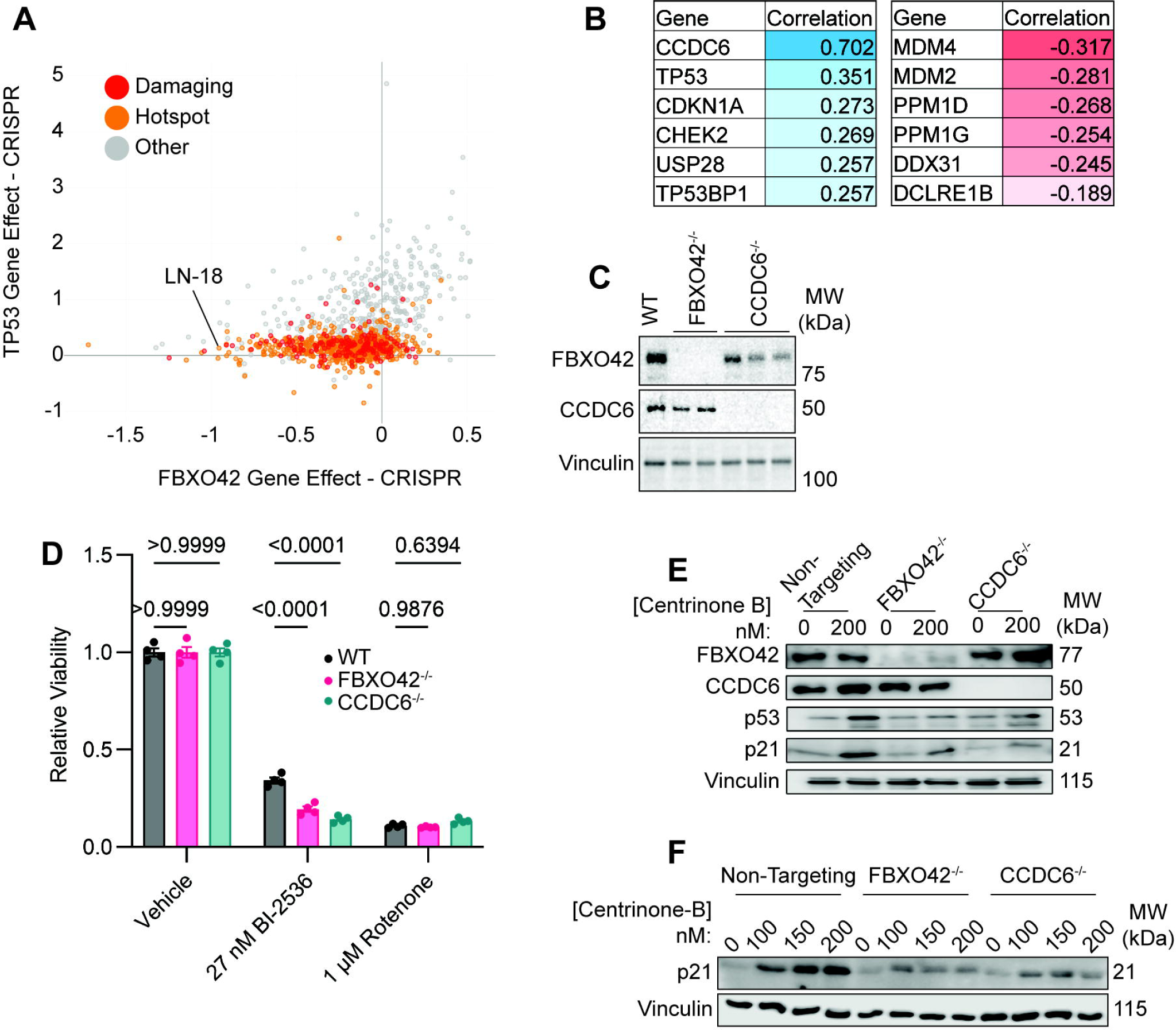
Involvement FBXO42 and CCDC6 in p53 Activation. **A.** DepMap data comparing the CRISPR essentiality scores of FBXO42 and p53 in the cell lines in the DepMap database. Colored dots indicate known p53 mutations in each cell line. LN-18 cells (see Figure 4) are highlighted. **B.** The top DepMap positive and negative co-dependencies of FBXO42. Values are Pearson correlations. **C.** Western blots of clonal RPE1 cell lines showing expression of FBXO42 and CCDC6. **D.** Presto blue-based viability assays of clonal RPE1 cell lines with the indicated genotypes. Cells were treated with BI-2536 (PLK1 inhibitor, 27 nM) or rotenone (electron transport chain inhibitor, 1 µM). Each point is a replicate well. Two-way ANOVA. p-value of individual comparisons, with multiple comparison correction, is displayed. **E.** Western blots of extracts from RPE1 cells with FBXO42 or CCDC6 knocked out, immunoblotted for the indicated proteins. Cells were untreated or treated with 200 nM centrinone-B (PLK4i) for 48 hours. **F.** As **E**, but cells were treated with 100, 150, or 200 nM centrinone-B.

### FBXO42 Ubiquitinates PPP4C

Since FBXO42 is a SCF E3 ligase substrate adaptor, it is likely that its functions in cell survival and mitosis are mediated by a substrate that FBXO42 binds and ubiquitinates.

To identify substrates of FBXO42, we performed immunoprecipitation on lysates from HAP1 cells of endogenous FBXO42 followed by mass spectrometry proteomics (IP-MS) (Figure 3A). Wild-type cells were compared with FBXO42^-/-^ cells. As expected, cullin-RING ligase complex members CUL1 and SKP1, and the suggested substrate RBPJ (Jiang et al., 2022), were identified as FBXO42 interactors. We also identified protein phosphatase complex 4 (PP4) members PPP4C, PPP4R1, and PPP4R2 (Figure 3A), as seen in previously published high-throughput immunoprecipitation mass spectrometry analysis (Huttlin et al., 2021; Jiang et al., 2022; Oughtred et al., 2021). We further analyzed FBXO42-CCDC6-PPP4C interactions by co-immunoprecipitation western blot, in RPE1 cells expressing epitope-tagged alleles of PPP4C, FBXO42, and CCDC6. Using co-immunoprecipitation, we observed that PPP4C interacts with both FBXO42 and CCDC6 (Figure 3B). PPP4C-FBXO42 interactions were maintained in CCDC6^-/-^ cells, and PPP4C-CCDC6 interactions were maintained in FBXO42^-/-^ cells (Figure 3B).

**Figure 3:**
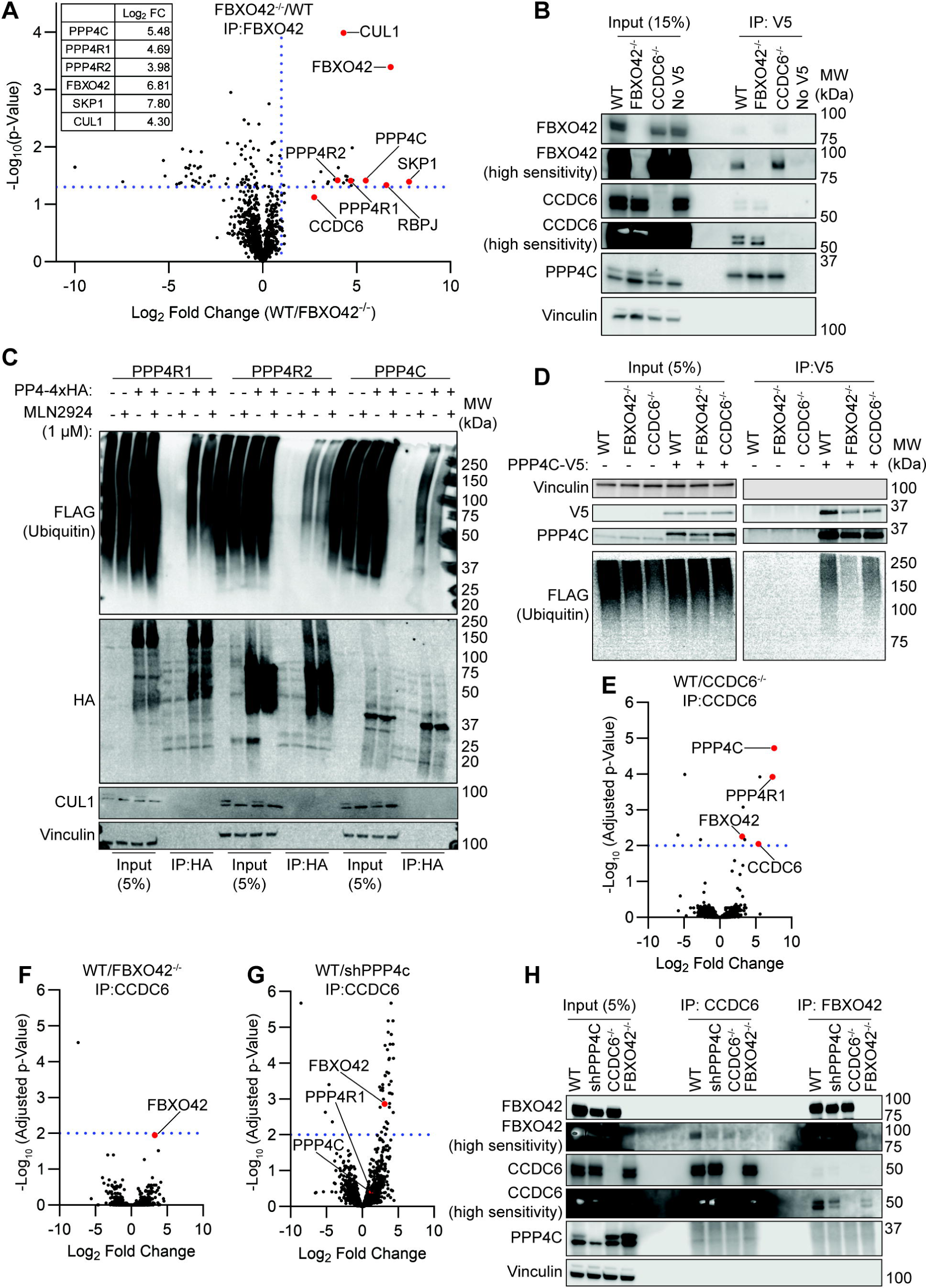
Interactions of FBXO42, CCDC6, and PPP4C. **A.** Immunoprecipitation-mass spectrometry (IP-MS) analysis of proteins interacting with FBXO42 in HAP1 cells. Enrichment of proteins in WT cells compared to FBXO42^-/-^ cells is shown. Fold change values for PP4 complex and SCF E3 ubiquitin ligase complex members are in the inset table. **B.** Immunoprecipitation (IP) western blotting performed on lysates from RPE1 cells. Expressing FBXO42-Myc, CCDC6-His, and where indicated PPP4C-V5. WT, FBXO42^-/-^, CCDC6^-/-^, or WT cells lacking PPP4C-V5 expression were used. FBXO42 and CCDC6 were knocked out, and clonal cell lines were used. Epitope tagged alleles are not present in the cell lines where the genes are knocked out or knocked down. 15% input compared to IP or immunoprecipitation with a V5 epitope tag targeting antibody is shown. High sensitivity panels show exposures of the same membranes developed with high sensitivity ECL reagent. **C.** IP western blotting of lysates from RPE1 cells. All cells express FLAG-tagged ubiquitin from a doxycycline (dox) induced promoter, induced with 1 µg/ml dox 24 hours before harvest. MLN4924 (pan-cullin-RING ligase inhibitor) added at 1 µM to indicated cells 24 hours before harvest. 10 µM MG-132 was added to cells 4 hours before harvest. Cells express PPP4R1-4xHA, PP4R2-4xHA, or PPP4C-4xHA as indicated. 5% Input compared to IP. IP was performed with beads with cross-linked anti-HA antibodies. CUL1 blot shows double bands in vehicle treated and a single band in MLN4924 treated lanes, consistent with inhibition of CUL1 neddylation. **D.** IP western blot of lysates from RPE1 cells. All cells express FLAG-tagged ubiquitin from a doxycycline (dox) induced promoter, induced with 1 µg/ml dox 24 hours before harvest. 10 µM MG-132 was added to cells 4 hours before harvest. Indicated cells express PPP4C with a V5 epitope tag. 5% Input compared to IP. IP was performed with a V5 antibody. **E-G.** IP-MS analysis. Lysates from RPE1 cells were immunoprecipitated with an anti- CCDC6 antibody. Comparison of lysates from WT cells to **E.** CCDC6^-/-^ cells. **F.** FBXO42^-/-^ cells. **G.** PPP4C knockdown cells. PPP4C was knocked down by lentivirus transduction with two shRNAs targeting PPP4C three days before cells were harvested. Proteins are expressed at endogenous levels. **H.** IP western blot. Similar to **B**, FBXO42-Myc, CCDC6-His, and PPP4C-V5 are expressed, except where they have been knocked out or knocked down. Immunoprecipitation of CCDC6 or FBXO42 performed with antibodies against those proteins. 5% input compared to IP. High sensitivity panels show exposures of the same membranes developed with high sensitivity ECL reagent.

To determine if FBXO42 ubiquitinates PP4 subunits, we generated a doxycycline (dox) induced, FLAG-tagged ubiquitin construct, and 4xHA tagged PPP4C, PPP4R1, and PPP4R2 constructs and introduced these into RPE1 cells. We immunoprecipitated HA and blotted for FLAG (ubiquitin) or HA (PP4 subunits). Cells were treated with a vehicle control or with MLN4924, which broadly inhibits cullin ring ligases (Baek et al., 2020; Brownell et al., 2010; Soucy et al., 2009). We observed a reduction in the polyubiquitin smear in MLN4924 treated cells with PPP4C, but not PPP4R1 or PPP4R2 (Figure 3C). We then tested ubiquitination of PPP4C in WT, FBXO42^-/-^, or CCDC6^-/-^ RPE1 cells. We immunoprecipitated V5-tagged PPP4C and blotted for FLAG-Ubiquitin. PPP4C showed a high-molecular weight smear, typical of ubiquitinated proteins. FBXO42 knockout cells displayed reduced ubiquitination of PPP4C (Figure 3D). In contrast, mutation of CCDC6 did not decrease PPP4C ubiquitination.

Since both FBXO42 and CCDC6 bound PPP4C, we explored whether the FBXO42- CCDC6 interaction was mediated by their simultaneous binding to PPP4C. We performed IP-MS analysis, using an antibody against endogenous CCDC6, with lysates from RPE1 cells that were wild type, FBXO42^-/-^, CCDC6^-/-^, or had PPP4C knocked down using simultaneous transduction of two shRNAs (shPPP4C) (Figure S3). Complete knockouts of PPP4C could not be generated, because PPP4C is essential in most, if not all, cells. Comparing WT cells to CCDC6^-/-^ control cells (Figure 3E), we observed enrichment of PPP4C, PPP4R1, and FBXO42, consistent with CCDC6 interacting with these proteins. In FBXO42^-/-^ cells (Figure 3F), there was no significant change in CCDC6 interactions other than with FBXO42 itself, suggesting that FBXO42 has little or no effect on CCDC6 interactions with the PP4 complex or any other proteins. Comparing WT cells to shPPP4C cells we observed that FBXO42 was enriched in WT cells over the shPPP4C cells, indicating that CCDC6-FBXO42 interactions were decreased when the levels of PPP4C were reduced (Figure 3G). We then examined whether the FBXO42-CCDC6 interaction depended upon PPP4C using co-immunoprecipitation western blotting. We immunoprecipitated either FBXO42 or CCDC6, and saw that in both cases knockdown of PPP4C decreased the FBXO42-CCDC6 co-immunoprecipitation to background levels (Figure 3H).

We used Alphafold 3 (Abramson et al., 2024; Jumper et al., 2021) to predict possible interactions between FBXO42, CCDC6, and PPP4C. Alphafold was not able to predict a complex of PPP4C and FBXO42 with adequate confidence. Alphafold did predict an interaction between PPP4C and CCDC6 (Figure S3B-G). CCDC6 has two predicted binding sites for PPP4C. The first predicted binding site showed hydrogen bonds between PPP4C and the CCDC6 residues Gln102, Glu110, Asn114, Lys118 (Figure S3B, S3D). When these residues are mutated to alanine, Alphafold predicts a second binding site, with hydrogen bonds between PPP4C and CCDC6 residues Glu 203, Glu211, Asn215, and Lys219 (Figure S3C). Notably, both binding sites contain the motif EXXXNXXXK. It is possible that this motif mediates PPP4C-CCDC6 interactions. CCDC6 has predicted disordered regions from residues 1-47 and most of 342-474 (the C-terminal) (Consortium, 2024). We repeated Alphafold predictions with just the CCDC6 residues 48-341 (Figure S3E-G). This generally increased the ipTM values of the predictions, and PPP4C still was predicted to interact with CCDC6 at the same residues.

### Knockdown of PPP4C Rescues Sensitivity to FBXO42/CCDC6 Knockdown in a Glioblastoma Cell Line

We hypothesized that FBXO42 and CCDC6 negatively regulate the activity of PP4, and that the loss of this regulation is responsible for their essentiality in some cell lines, such as the glioblastoma line LN-18 (Figure 2A). We analyzed viability and growth of LN-18 cells transduced with shRNA against FBXO42 or CCDC6 using a flow-cytometry-based competitive growth assay. As expected, we observed that FBXO42 or CCDC6 knockdown reduced cell fitness compared to scramble controls (Figure 4A-B). Knockdown of FBXO42 increased protein levels of PPP4C (Figure 4C-D, S4A). To test if increased levels or activity of PPP4C contributed to the reduced proliferation of FBXO42 or CCDC6 knockdown cells, we simultaneously knocked down PPP4C and either FBXO42 or CCDC6, where the shRNA against FBXO42, CCDC6, or scramble control where in a vector with a GFP marker, and each of the three anti-PPP4C shRNA, and their scramble control, were in a GFP-negative vector with a puromycin resistance marker (Puro). We were unable to generate stable PPP4C knockout clones, since it is an essential gene. Cell viability was measured using Presto Blue. Growth for the Puro vector shRNAs were normalized to that of cells with that GFP scramble control (Figure 4E). In parallel experiments, we compared the growth of the FBXO42 or CCDC6 knockdown with or without PPP4C knockdown in a competitive growth assay (Fig S4B). In both assays, we found that knockdown of PPP4C partially rescued the effects of either FBXO42 or CCDC6 knockdown, suggesting that these proteins negatively regulate PP4 activity in cells.

**Figure 4:**
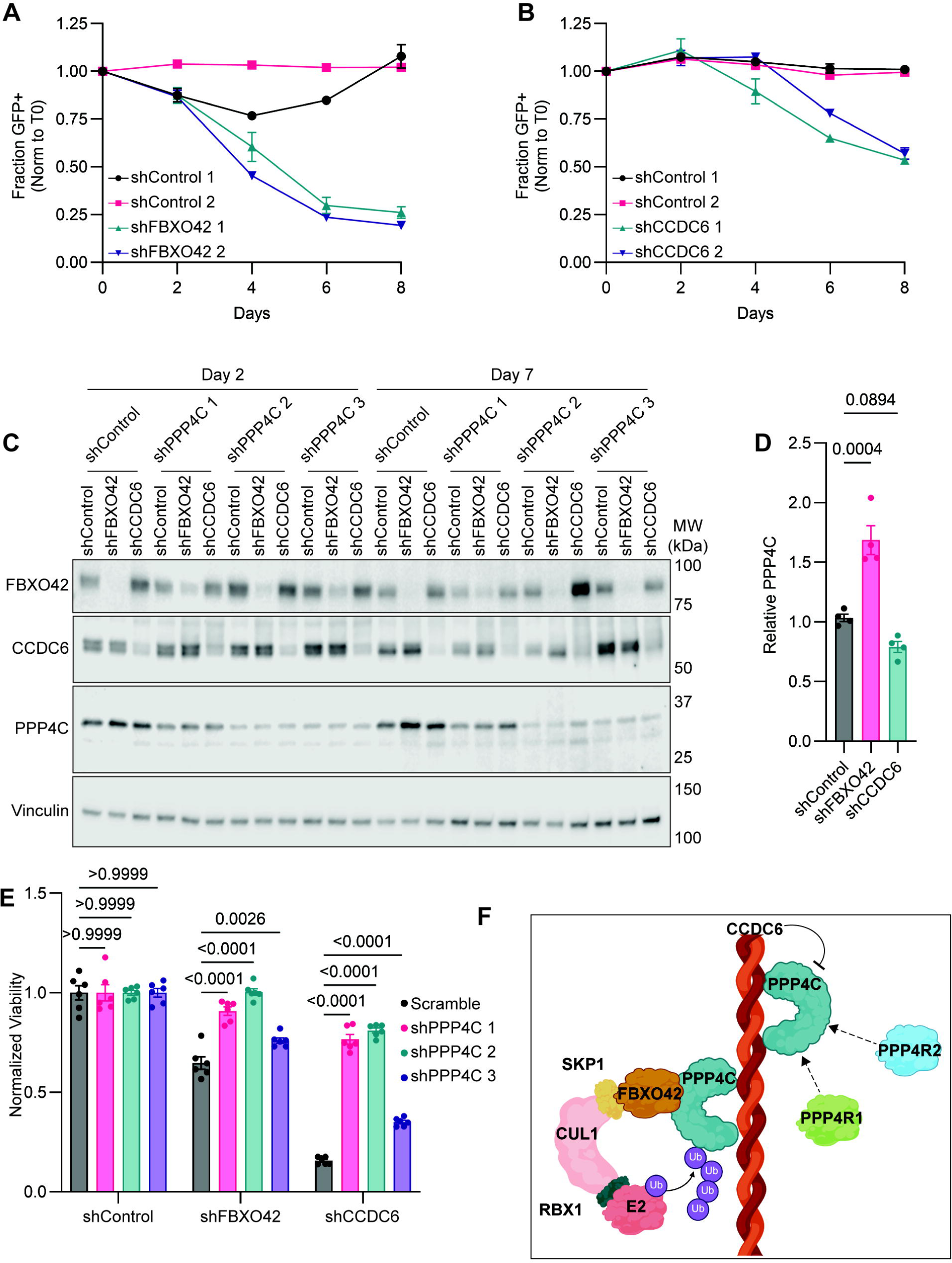
Knockdown of PPP4C Rescues Sensitivity to FBXO42/CCDC6 Knockdown in a Glioblastoma Cell Line. **A.** GFP, flow cytometry-based, competitive growth assay. LN-18 cells transduced using lentivirus with shRNA constructs against non-targeting sequences, or FBXO42. Each point represents a replicate well. Representative of three independent experiments. **B.** As **A**, but with shRNAs targeting CCDC6. **C.** Western blot analysis of LN-18 cells transduced with shRNA against PPP4C or a scramble control. Cells were also, simultaneously, transduced with shRNA targeting FBXO42, CCDC6, or a scramble control. **D.** Quantification of PPP4C protein levels from **Figure S4A**. Normalized to vinculin expression. One-way ANOVA, p-value of individual comparisons is displayed. Each point represents a replicate, parallel transduction and western blot lane. **E.** Presto Blue based viability assay. LN-18 cells were transduced with shRNA against FBXO42, CCDC6, or a scramble control using lentivirus. Cells were additionally transduced with shRNA against one of three shRNA sequences targeting PPP4C, or a scramble control. Equal numbers of cells were seeded, and viability was assessed six days after transduction. Viability was normalized to the scramble control paired with the FBXO42 or CCDC6 constructs. Two-way ANOVA, p-value of individual comparisons is displayed. Representative of 3 independent experiments. **F.** Schematic representing our hypothesized model of FBXO42-PPP4C-CCDC6 interaction. FBXO42 acts as a substrate adapter for PPP4C ubiquitination. Separately, PPP4C binds CCDC6, at two sites, which inhibits PPP4C activity. PPP4R1 and PPP4R2 were identified as interactors by IP-MS, but their interactions and roles are still unclear.

## DISCUSSION

Multiple recent studies have highlighted the importance of FBXO42 in mitosis and cancer (Barbosa et al., 2020; Hoellerbauer et al., 2024; Hundley et al., 2021; Jiang et al., 2022; Lü et al., 2024; Nagler et al., 2020; Zhou et al., 2024). However, a detailed understanding of its function, substrates, and contribution to cancer cell survival was lacking. Furthermore, the literature suggests that FBXO42 may work with CCDC6 in some fashion (Hoellerbauer et al., 2024; Lü et al., 2024), but the nature and function of their relationship was not well-understood. Here, we present data showing that CCDC6 and FBXO42 inhibit PPP4C in different ways: FBXO42 by targeting PPP4C for turnover, and CCDC6 by an as of yet understood, but previously characterized, mechanism (Morra et al., 2022).

The synthetic lethality between the γ-tubulin ring complex proteins MZT1 and MZT2B and FBXO42 mutation is consistent with our previous observation that FBXO42 knockout leads to the accumulation of cells with monopolar spindles (Hoellerbauer et al., 2024; Hundley et al., 2021). FBXO42 partially ameliorated reduced cell fitness caused by TUBE1 knockout. This might be caused by a reduction in p53 signaling, consistent with the finding that TUBE1^-/-^ cells arrest in a p53-dependent manner (Wang et al., 2017). The cell line dependencies of FBXO42 vs p53 (Figure 2A) show that cell lines in the upper right quadrant, whose growth rate increases with p53 mutation, also have increased growth rates when FBXO42 (or CCDC6) is mutated. However, unlike p53, which is almost never required for proliferation, many cell lines (enriched for those with mutated p53 pathways) actually require FBXO42 (and CCDC6) for viability. We found that FBXO42 binds to and ubiquitinates PPP4C in a CCDC6-independent manner and suppresses PPP4C expression. We also found that PPP4C contributes to cell death in FBXO42 knockout LN-18 cells. CCDC6 also interacts with PPP4C, and PPP4C knockdown also partially rescued cell fitness defects in CCDC6 knockout LN-18 cells. Given that over activation of PP4 can inhibit cell proliferation (Huang et al., 2016), we hypothesize that overactive PP4 causes cell death in FBXO42 and CCDC6 knockout cells. Cell lines such as LN-18 may possess intrinsic mitotic defects or have increased reliance on a kinase opposed by PP4.

CCDC6 binds to PP4 (Ueki et al., 2019), and reduces its activity (Morra et al., 2022), and we show that this interaction is independent of FBXO42. While CCDC6 has been reported to associate with PP4 through the PPP4R3 subunits (Ueki et al., 2019), we find that PPP4R1 co-immunoprecipitates with CCDC6. Since PPP4R1 and PPP4R3 subunits are in separate PP4 complexes, CCDC6 must recognize the PP4 complexes by at least one other mechanism. Similarly, FBXO42 immunoprecipitates not only PPP4C, but also PPP4R1 and PPP4R2 (Figure 3A), consistent with a model in which PPP4C is recognized directly by FBXO42. PPP4C forms complexes with PP4 regulatory subunits in different configurations (Park and Lee, 2020), and future investigation should determine to what extent FBXO42 and CCDC6 bind to and regulate these different complexes. FBXO42 has been shown to regulate synaptonemal complex assembly during meiosis in Drosophila by downregulating protein levels of PP2A-B56 (Barbosa et al., 2020). PP2A-B56 is a regulatory subunit of PP2A. PP2A and PP4 are closely related and evolutionarily conserved (Chen et al., 2017), so it is plausible that FBXO42 could regulate both complexes in both organisms. Alphafold predictions were not able to predict a PPP4C- FBXO42 interaction, but did predict a robust interaction between PPP4C and a repeating motif EXXXNXXXK that occurs twice within the CCDC6 coiled coils. Together our data suggest a model in which FBXO42 and CCDC6 bind and inhibit PPP4C independently (Figure 4F).

It remains unknown whether FBXO42’s regulation of PP4 accounts for its role in p53 signaling and mitosis. While reducing PPP4C expression partially rescued cell fitness in FBXO42 knockdown cells, this does not rule out the contribution of other FBXO42 substrates. Furthermore, it is unknown which PP4 substrates contribute to the phenotype of FBXO42^-/-^ cells, and whether it is the same substrate affecting both p53 signaling and mitosis. It has been shown that PP4 regulates microtubule organization via dephosphorylation of NDEL1 (Toyo-oka et al., 2008). We speculate that perturbation of this regulatory mechanism could account for the dysregulation of mitotic spindle assembly observed in FBXO42 knockout cells. PP4 also antagonizes p53 signaling (Shaltiel et al., 2014). Further study is also needed to elucidate the regulation of PPP4C targeting by FBXO42. PPP4C is not a highly unstable protein (Mathieson et al., 2018), suggesting that FBXO42 may only target a subset of it, or may target bulk levels in a subset of cells. Additionally, further study is needed to determine the mechanism of CCDC6 regulation of PP4. Answering these questions may reveal why FBXO42 is essential in only a subset of cancer cells, which could guide the development of new, targeted cancer therapies (Fischer et al., 2014; Senft et al., 2018).

## Supplemental Figure Legends

**Figure S2: Sequences of FBXO42 and CCDC6 knockout clones.**

Sanger sequencing results are shown for **A.** FBXO42 and **B.** CCDC6 knockout clones, RPE1 cells. Consensus coding sequences (CCDS) are shown for each gene, as is the expected binding site of the sgRNA used to generate knockouts. WT and mutated sequences are aligned with amino acid translations. Each knockout cell line carries a mutation generating a frame shift.

**Figure S3: Interactions of FBXO42, CCDC6, and PPP4C Continued**

**A.** Western blot showing expression of FBXO42, CCDC6, PPP4C, or vinculin (loading control) in the RPE1 cell lines used in Figure 3E-G.

**B-G.** Alphafold 3 predictions of CCDC6-PPP4C interactions. ipTM and pTM values shown. Protein representations colored by pIDDT values. **B-D.** Uses full length CCDC6, **E-G.** uses CCDC6 with its terminal disordered regions truncated, residues 48-341. **B and E** use WT CCDC6. In **C and F, and D and G** the indicated residues on CDC6 are mutated to alanine to ablate the two different predicted binding sites.

**Figure S4: Knockdown of PPP4C Rescues Sensitivity to FBXO42/CCDC6 Knockdown in a Glioblastoma Cell Line Continued**

**A.** Western blot showing expression of indicated proteins in LN-18 cells transduced with a scramble control, or shRNA against FBXO42 or CCDC6. Four replicate, parallel transductions were performed for each condition. All panels for each protein are from the same image with the same brightness and contrast settings, but with unnecessary lanes cropped out.

**B.** GFP-based competitive growth assay analysis of LN-18 cells treated similarly as in Figure 4D. Fraction of GFP expressing cells are shown, normalized to time zero, and to the Scramble-GFP control. Analyzed on day 6 post transduction.

## METHODS

### Human Cell Culture

All cells were cultured in a humidified, 37 °C incubator, with 5% CO2 atmosphere. 10% heat inactivated fetal bovine serum (FBS), 10 mM HEPES pH 7.5, and 1X Antibiotic-Antimycotic (Anti-Anti) was added to all media unless otherwise specified. RPE1 cells were cultured in DMEM/F12 media (Thermo Fisher Scientific 11-320-033). LN-18 cells and HEK 293T cells were cultured in DMEM, high glucose, no glutamine (Thermo Fisher Scientific 11960069), with 1X GlutaMAX added. HAP1 cells were cultured in IMDM (Thermo Fisher Scientific 12440061) without added HEPES. DPBS without calcium and magnesium were used to wash cells, Trypsin-EDTA (0.25%) was used to dissociate cells. Cells were maintained in aseptic conditions using standard culture methods, with subculturing every 3 or 4 days at split ratios ranging from 1:10 to 1:20 depending on the growth of the cells.

### CRISPR Screen

The production of the lentivirus and HAP1 Cas-9 clones have previously been described (Hundley et al., 2021; Hundley and Toczyski, 2021). The human GeCKO v.2 library (Sanjana et al., 2014) was transduced into HAP1-Cas9 cells or clonal HAP1-Cas9 cell line harboring an FBXO42 deletion. 24 hours after transduction, lentivirus containing media was replaced with fresh IMDM. 24 hours after media replacement, HAP1 cells were split into two separate populations (replicate 1 and 2) and treated with 1 μg/mL puromycin and 1 μg/mL of doxycycline to select for sgRNA integration and initiate Cas9 production, respectively. Cells were maintained at 1,000x coverage. After 3 days of puromycin selection and doxycycline induction approximately 150x10^6^ cells were frozen for the T0 time point. Cells were then split into three different conditions: untreated group, 3nM of Bi-2536 or 10nM colchicine for 8 days being split every 2 days. Cells were maintained at a coverage >600x. After 8 days of treatment, cells were collected with each replicate having >150x10^6^ cells.

### Next generation sequencing library preparation

Genomic DNA was extracted from cell pellets using Nucleospin blood XL midi kit (Machery Nagel). The sgRNA-encoding regions were amplified by PCR with primers containing Illumnia sequencing adaptors and barcodes. Each 100ul PCR contained 10ug of DNA. One PCR contained no gDNA to ensure there was no background amplification. 5 µl of each PCR was run on an agarose gel, and all the samples were pooled by their relative intensities. The pooled sample was then gel purified and run on a HiSeq 4000 with single end, 50 base reads (UCSF Center for Advanced Technology) using common 3′ and 5′sequencing primers and an indexing primer provided by the core facility (Hundley et al., 2020). Reads were trimmed and aligned following previously published methods (Horlbeck et al., 2018).

### Crystal Violet Assay

A different WT and FBXO42^-/-^ HAP1-Cas9 clone was used to validate MZT1, MZT2B and TUBE1 as hits from the screen. To validate these hits, individual sgRNAs—including a guide targeting a non-expressed gene (non-targeting guide)—were cloned and transduced into each cell line. The cells were then treated with 1μg/mL puromycin and 1μg/mL of doxycycline to select for sgRNA integration and initiate Cas9 production, respectively. After three days of selection and Cas9 expression, 1000 cells were seeded into a 6-well dish in IMDM media, or IMDM media with 3 nM Bi-2536. Media was changed every 3-4 days for a 10 day incubation. After 10 days, cells were washed with PBS and stained with crystal violet following the Xin Chen Lab’s protocol (Chen) (https://pharm.ucsf.edu/xinchen/protocols/cell-enumeration). In short, cells were stained with crystal violet, washed with PBS and lysed in 500 uL of lysing solution. The experimental lysate was diluted 1:10 in PBS and the OD was measured at 590nm.

### Establishment of Clonal RPE1 FBXO42 and CCDC6 Knockout Lines

Knockouts were generated through electroporation-based transfection of Cas9-sgRNA ribonucleoprotein particles. The sgRNA sequences (sequence of DNA targeted) were: CCDC6: GCTCTCCAGAAAATTGATGC, FBXO42: AGCATGGGGCCCCAGAACTG. The Neon Electroporation system was assembled in a sterile biosafety cabinet according to manufacturer instructions. Work was all performed under sterile conditions, and the work area, pipettes, and gloves of researchers were cleaned with RNAse-Zap. Purified sgRNA were ordered from Thermo Fisher Scientific. sgRNA was resuspended at 100 µM in sterile, RNAase-free TE buffer (10 mM Tris-HCl pH 8.0, 0.1 mM EDTA). Purified Truecut Cas9 Protein (Thermo Fisher), supplied was supplied at 5 µg/µl. For each reaction, 2 µg (0.4 µl) Cas9 was combined with 12 pmol sgRNA, brought to a total volume of 5 µl with Buffer R (Thermo fisher). We have observed that PBS without Calcium and Magnesium is an acceptable substitute for Buffer R. This mixture for a single reaction, was scaled up to perform 6 reactions per sgRNA construct. This mixture was incubated for at least 10 minutes at room temperature to allow for complex formation.

A twelve well plate was prepared with antibiotic-free media, and placed in the 37°C, 5% CO2 incubator to equilibrate. Cells were washed with DPBS, and trypsinized. Trypsin was quenched with media containing 10% FBS. Cells were counted using a hemocytometer or an automated cell counter. Cells were centrifuged (300 g, 5 minutes) and washed with DPBS. Cells were centrifuged again, and resuspended in Buffer R at a concentration of 25 million viable cells per ml. Cells were mixed at a 1:1 ratio to the pre-complexed Cas9- sgRNA, and mixed gently, taking care to avoid bubble formation. Electroporation was performed according to manufacturer instructions, with the electroporation conditions of 1300V, 2 pulses, 20 ms pulse width. Electroporated cells were deposited in a 12-well plate well with pre-warmed, antibiotic free media, one well per electroporation (approximately 125,000 cells/well).

Two days later, cells were passaged, seeded for single cell, and some were collected for analysis of knockout efficency. Approximately 100,000 cells were collected, and DNA was extracted using QuickExtract according to manufacturer instructions. Extracted genomic DNA was PCR amplified and sent for Sanger sequencing. ICE analysis (Conant et al., 2022) was used to estimate the fraction of cells with InDels and knockout inducing mutations. This information was used to estimate how many clones should be screened for knockout mutations. A FACS instrument was used to deposit single cells into wells of a 96 well plate to isolate clones. Single cells grew for approximately two weeks until there were enough cells to analyze, and then clones were sequenced using sanger sequencing or the region flanking the expected CRISPR cut sites. The following primers were used for sanger sequencing and ICE Analysis. For FBXO42, PCR Forward: GGCCCCTGTAAGTTTTCCCA, PCR Reverse: CCATGTGCTAAGCCAGTGGTA, Sanger Sequencing: CTTTCAAGACCAGACAGGAGAG; CCDC6: PCR Forward: AGCAAAGAATAAAGAGGTGCCC, PCR Reverse: ACTAGTACACCGAACTCCCC, Sanger Sequencing: GGCCACATGTTTGCAGTATTC. Clones with predicted knockouts (e.g. mutations causing a frameshift, see Figure S2) were further validated using western blotting.

### Lentivirus Production and Transduction

All handling of lentivirus was performed with BSL-2+ safety procedures. Lentivirus packaging was performed with 2^nd^ generation lentiviral packaging plasmids psPAX2 (a gift from Didier Trono (Addgene plasmid # 12260; http://n2t.net/addgene:12260; RRID:Addgene_12260)) and pMD2.G (a gift from Didier Trono (Addgene plasmid # 12259; http://n2t.net/addgene:12259; RRID:Addgene_12259)). For lentivirus production, 9.5 x 10^6^ HEK 293T cells were seeded in one T-75 flask per lentiviral construct, in 16 ml DMEM based culture media. The next day, media was aspirated and replaced with 8 ml lentivirus packaging media (Opti-MEM, 5% FBS, 200 μM sodium pyruvate, no antibiotics, no added HEPES). 800 μl Opti-MEM, 1940 ng pMD2.G, 2656 ng psPAX.2, and 3978 ng of the transfer plasmid containing the gene or shRNA sequence of interest were combined and incubated for 10 minutes at room temperature with periodic mixing. This mixture was added to the cells. Four hours later, media was changed to 16 ml fresh lentivirus packaging media. Virus containing media was collected 24 and 48 hours after transfection and combined. Virus containing media was centrifuged at 150 x g for seven minutes to pellet any loose HEK 293T cells. The virus-containing supernatant was then filtered with a 0.45 μm syringe filter. Transfections could be scaled up or down, depending on how much virus was required, based on the surface area of the cultureware used. Virus containing media was stored at 4 °C for up to about 2 months, or snap frozen and stored at -80 °C.

For transductions, cells were passaged the day before such that they were at about 20% confluence on the day of transduction. Cell media was replaced with fresh, complete culture media with 8 μg/ml polybrene. Virus containing media was then added at a dilution factor determined by empirical optimization, typically 1:8 or 1:16. Media containing virus was removed the following day and replaced with fresh culture media.

### Plasmids

GFP expressing shRNA Constructs were cloned into a plasmid backbone where EGFP is expressed from a hPGK promoter, and shRNA sequences are expressed from a U6 promoter. shRNA targeting sequences were: FBXO42 1: CAAACAGTGGTATCGACTTAT, FBXO42 2: TAGATGATGCAACTATCTTAA, CCDC6: GCCTACTCCTTCGCAACATTC, Scramble control: CCTAAGGTTAAGTCGCCCTCG.

Plasmids expressing shRNA against PPP4C were purchased from the Sigma Aldrich MISSION shRNA Collection, and are in the pLKO.1 vector. The following are the TRC IDs of the constructs, followed by the targeted sequence: shPPP4C 1: TRCN0000002761, CGGCTACCTATTTGGCAGTGA; shPPP4C 2: TRCN0000002763, CATCAAGGAGAGCGAAGTCAA; shPPP4C 3: TRCN0000010737, GTGCTTCGAGGGTGAGGACTT. The scramble control, in the same vector, was acquired from Addgene, a gift from Anthony Leung (Addgene plasmid # 136035; http://n2t.net/addgene:136035; RRID:Addgene_136035).

For the plasmid expressing FLAG-tagged ubiquitin, the pLVX-TetOne backbone was used. Backbone was linearized with EcoRI and BamHI restriction enzymes. Insert contains a FLAG tag, the coding sequence for ribosomal protein RPS27A (which includes an N-terminal fusion with ubiquitin, that is cleaved off to form free ubiquitin post-translationally), followed by a “self cleaving” P2A sequence, then EGFP. Inserts were PCR amplified and the whole construct was assembled by Gibson assembly (Gibson et al., 2009). After this construct was transduced into RPE1 cells (see above), GFP expression was induced by doxycycline treatment (1 µg/ml, 24 hours) and cells were sorted by FACS to select for GFP expressing cells. FACS selection was repeated three times to select for cells that consistently express the FLAG ubiquitin containing plasmid. Cell GFP expression (following dox induction) was periodically monitored to ensure the cells maintained consistent expression.

To overexpress epitope tagged versions of PP4 subunits, cDNA sequences for PPP4C, PPP4R1, and PPP4R2 in pLenti6.3 were purchased from DNAsu. The V5 tag was replaced by 4 x HA tags using a gBlock, PCR amplification of the backbone, and Gibson assembly. Constructs were transduced into RPE1 cells with lentivirus (see above), and cells were selected with 5 μg/ml blasticidin.

To overexpress epitope-tagged versions of CCDC6 and FBXO42, the protein coding sequences were obtained as gBlocks (IDT). They were cloned into a pLVX-TetOne-Puro backbone using Gibson cloning. In both cases a GGGS linker was added after the c-terminal of the gene before the epitope tag. A 6xHis tag (six histidines) was added to CCDC6, and a Myc tag (EQKLISEED) was added to FBXO42. Linkers and tags were added by PCR using the following primers: FBXO42-Myc forward: ctgattagcgaagaagatctgTAAGGATCCAGACCACCTC, FBXO42-Myc Reverse: tttctgttcagagcctccaccTCTCTTTGCTCGTACAAAG, CCDC6-His Forward: catcatcaccaccacTAAGGATCCAGACCACCTCCC, CCDC6-His Reverse atgagagcctccaccAGGCTGGGAGGAGGGGTG.

Constructs were transduced into RPE1 cells with lentivirus (see above), and cells were selected with 5 μg/ml puromycin. We made versions of the plasmids where the Puro selection marker was replaced by TurboRFP. TurboRFP was amplified from pCW57-RFP- 2A-MCS (gift from Adam Karpf (Addgene plasmid # 78933; http://n2t.net/addgene:78933; RRID:Addgene_78933)) with these primers. Forward: GTTGATTGTTCCAGACGCGTATGAGCGAGCTGATCAAGGA, Reverse: GGCTTTTGCAAAACGCGACCTTATCTGTGCCCCAGTTTGC. The plasmid backbones containing FBXO42 or CCDC6 coding sequences were amplified with the following primers, which flank the area around the Puro gene. Forward: GCAAACTGGGGCACAGATAAGGTCGCGTTTTGCAAAAGCC, Reverse: TCCTTGATCAGCTCGCTCATACGCGTCTGGAACAATCAACC. PCR reactions were performed using SuperFI Platinum Taq II according to manufacturer instructions. Amplified backbones were digested with DpnI to remove parental plasmids, prior to Gibson assembly. Constructs were transduced into RPE1 cells with lentivirus (see above), and cells were selected with at least three rounds of sorting based on TurboRFP expression.

### Presto Blue Viability Assay

LN-18 cells were seeded to 96-well plates at 3000 cells per well, with 200 µl culture media per well. For experiments with BI-2536 and rotenone, cells were seeded to 96 well plates in 100 μl media. The following day, compounds were added to wells at 2 times the desired final concentration, in 100 μl per well. For viability analysis, Presto Blue was used according to manufacturer instructions with the following modifications. Presto Blue concentrate was diluted 1 part to 10 parts complete, 37°C culture media. Culture media was removed from the cells by firmly flicking the plate over a large beaker. Residual media was removed by tapping the plate, inverted, on a lint-free paper wipe (e.g. Kimwipe). 100 µl presto blue in media was added to each well. The plate was incubated at 37°C for 1 hour. Plates were then analyzed with an Agilent BioTek Synergy Neo2 plate reader using fluorescent mode, excitation 560 nm, emission 590 nm. On each plate, wells with no cells were given media and other manipulations and used as blanks. The average blank signal was subtracted from all other well signals during analysis.

### Western Blots

Cells were lysed by resuspending pellets in NP-40 based lysis buffer (100mM Tris pH 8, 150mM NaCl, 5mM EDTA, 0.1% NP-40, complete Protease Inhibitor tablet, and PhoSTOP tablet 100 mM PMSF added right before use). Cells were incubated at 4°C on a rotator for 30 minutes. Lysates were clarified by centrifuging at 20,000 x g for 20 minutes, and supernatants were transferred to new tubes. Protein concentrations were measured using the bicinchoninic acid (BCA) assay (Smith et al., 1985). Up to 30 μg protein was loaded per SDS-PAGE gel lane. Western blotting was performed using standard methods (Burnette, 1981). Proteins were resolved using 4-15% or 4-20% polyacrylamide Bio-Rad Criterion TGX Midi gels, with tris-glycine-SDS running buffer, at 150 V for approximately 90 minutes. Gels were transferred to nitrocellulose membranes using a “wet transfer” method, with a BioRad Criterion Blotter, and transfer buffer with 20 mM Tris, 150 mM glycine, 0.08% SDS, and 20% methanol, at 100 V for 60 minutes, at 4°C. Ponceau stains were performed to verify correct transfer. Blocking was performed using blocking buffer according to the primary antibody manufacturer instructions, usually 5% (w/v) bovine serum albumin (BSA) or non-fat dry milk in PBS with 0.05% tween (PBST). Importantly, we found that when blotting with the FBXO42 antibody, 3% non-fat dry milk in PBS with tween produced the best results. Primary antibodies were diluted according to manufacturer instructions in blocking buffer, typically at 1:1000. Sodium azide was added to the blocking buffer at 0.02% to prevent microbial growth. Blots were incubated with primary antibody overnight at 4°C on an orbital shaker, overnight. Primary antibody dilutions could typically be reused multiple times. Following primary antibody incubation, blots were washed 3-4 times with PBST for five minutes each. HRP conjugated secondary antibodies were diluted 1:10,000 in the same blocking buffer as the primary antibody (without sodium azide). Blots were incubated with diluted secondary antibody for 1 hour at room temperature, on an orbital shaker. For imaging, Pierce ECL Western Blotting Substrate was used as a substrate for chemiluminescence. For less abundant proteins in immunoprecipitation experiments, membranes were subsequently imaged again with SuperSignal West Femto Maximum Sensitivity Substrate (images marked as “high sensitivity” in figures). Imaging was performed using a LiCor XF.

### Immunoprecipitation

For immunoprecipitation experiments, all cells were treated with 10 μM MG-132 for four hours before collection. Cells were washed with DPBS and rapidly frozen on dry ice, as dry pellets. Pellets were lysed in lysis buffer (100 mM Tris pH 8, 150 mM NaCl, 5% Glycerol, 5 mM EDTA, 0.1% NP-40, complete Protease Inhibitor tablet, and PhoSTOP tablet) and passed through a 21g needle ten times. Lysates were rotated at 4 °C for 30 minutes. Lysates were then cleared at 20,000 x g for 10 minutes at 4 °C. Supernatant was saved and a BCA assay was performed to measure protein concentration. At least 900 μg total protein lysate was used for each immunoprecipitation. Antibody was added at 9 μg per condition, in 300 μl lysis buffer, and lysates were then incubated at 4C for 2 hours with rotation. Protein A Dynabeads were pre-washed three times in 1ml of lysis buffer and then resuspended in the original volume with lysis buffer. 300 µl samples were then incubated with 150 µl of Protein A Dynabeads for 1 hour at 4 °C with rotation. After the bead incubation, flow through was saved before the beads were washed five times with 1 mL of lysis buffer. At each wash step, beads were mixed by pipetting. To elute, Elution Buffer (50 mM Tris pH 8.0, 0.5% SDS, 12.5 mM EDTA) was added to each sample and samples were incubated at 65C for 10min. The elution was saved and frozen at -80 °C for downstream analysis.

The FBXO42 IP-MS was performed using HAP1 WT cells and HAP1 FBXO42^-/-^ clones, described in (Hundley et al., 2021), in duplicate. Cells were cultured in IMDM media supplemented with 10% FBS and scaled up to six 15 cm dishes. Cells were treated with 10 μM MG132 being added in the last 4 hours of treatment. The CCDC6 IP-MS was performed the same way, except using RPE1 cells. For the shPPP4C condition, RPE1 cells were transduced with PPP4C shRNA 1 and 2 (see above) in equal parts, 3 days before harvest. For experiments with overexpressed FBXO42, CCDC6, and PPP4C, expression of FBXO42 and CCDC6 was induced by adding doxycycline to the cell culture media at 2 μg/ml, 2 days before cell collection.

For ubiquitination experiments, FLAG-ubiquitin expression was induced by addition of doxycycline to the cell culture media at 1 μg/ml 24 hours before cell collection. MLN4924 was added to cells at 1 μM final concentration 24 hours before harvest. For these experiments, 25 mM N-Ethylmaleamide (NEM) was added to the lysis buffer, as a DUB inhibitor. Immunoprecipitation was performed with anti-HA or anti-V5 beads according to manufacturer instructions. Washes were performed using RIPA buffer (50 mM Tris-HCl pH 7.4, 150 mM NaCl, 1% Triton X-100, 0.5% sodium deoxycholate, 0.1% SDS, 1 mM EDTA) in order to remove non-covalent protein-protein interactions, leaving covalent ubiquitin-substrate interactions.

### Mass-spectrometry proteomics analysis

The raw data files were analyzed using FragPipe to obtain protein identifications and their respective label-free quantification values. Of note, the data were normalized based on the assumption that the majority of proteins do not change between the different conditions. Data was first filtered for contaminant proteins. In addition, proteins that have been only identified by a single peptide and proteins not identified/quantified consistently in the same condition have been removed as well. The data was converted to log_2_ scale, samples were grouped by conditions and missing values were imputed using man method, which uses random draws from a left-shifted Gaussian distribution of 1.8 StDev (standard deviation) apart with a width of 0.3. Protein-wise linear models combined with empirical Bayes statistics were used for the differential expression analyses. The limma package from R Bioconductor was used to generate a list of differentially expressed proteins for each pair-wise comparison. A cutoff of the adjusted p-value of 0.05 (params$fdr_correction method) along with a |log2 fold change| of 1 has been applied to determine differentially expressed proteins in each pairwise comparison.

### Flow Cytometry (Competition Assay)

For flow-cytometry, GFP based, competitive growth assays, LN-18 cells were transduced using high titer lentivirus to express shRNA constructs with GFP markers. The next day, cells were dissociated and analyzed by flow cytometry to ensure high GFP expression. Cells were mixed at an approximately 1 to 1 ratio with untreated, GFP negative cells. The mixed cells were then analyzed by flow cytometry to establish a time zero GFP expression fraction. Mixed GFP+ and GFP- cells were then seeded to 96 well plates at 3000 cells per well. At various time points, cells were analyzed by flow cytometry using a modified version of established protocol (Spangenberg et al., 2021). Media was removed from plates by flicking (in a biosafety cabinet to contain potentially hazardous aerosols). Cells were washed with DPBS (again removed by flicking). 25 μl of 0.25% trypsin was added to each well, and plates were incubated at 37 °C until cells appeared loosened under the microscope. Then, 200 μl cold FACS Buffer (DPBS with 2% FBS, 1 mM EDTA, 10 mM HEPES and (optionally) 1X Antibiotic-Antimycotic) was added to each well. The cells in each well were mixed by repeated pipetting. Cells were kept at 4 °C or on ice from this point on, until analysis. An Attune NxT flow cytometer was used to analyze cells and establish the percent GFP+.

For combined shRNA experiments, cells were transduced with GFP marked shRNA constructs (targeting FBXO42, CCDC6, or scramble controls) and puromycin resistance (Puro) marked shRNA constructs (targeting PPP4C or a scramble control) at the same time. The next day, media was removed, and fresh media with 2 μg/ml puromycin was added. After 3 days of puromycin selection (and four days post-transduction) cells were dissociated, analyzed, and mixed as described above.

For presto blue analysis, cells were treated and set up in the same way, except that instead of mixing GFP+ and GFP- cells, just 3000 GFP+ cells were seeded to each 96- well plate well.

### Alphafold Predictions

Performed with Alphafold 3 Server. Amino acid sequences for CCDC6 (Q16204) and PPP4C (P60510) were retrieved from UniProt.

## ACKNOWLEDGEMENTS

This work was supported by NIH grants R35 GM118104 to D.P.T., and R35 GM153408 to J.A.W., funding from the UCSF RAP Program to D.P.T., the UCSF Program for Breakthrough Biomedical Research (funded in part by the Sandler Foundation) to D.P.T., the UCSF Center for Advanced Technology, and the UCSF Laboratory for Cell Analysis which is supported by the NCI grant P30CA082103. The Gilbert lab was funded by the Arc Institute, NIH DP2CA239597, and a Pew-Stewart Scholars for Cancer Research award. S.H.S. is supported by the NCI Training Grant T32CA108462.

## Notes

### Competing Interest Statement

The authors have declared no competing interest.

