## Supplementary figures and images for "The regulation of Protein Phosphatase 4 by FBXO42 is required for cancer cell survival"

### Supplemental Figure

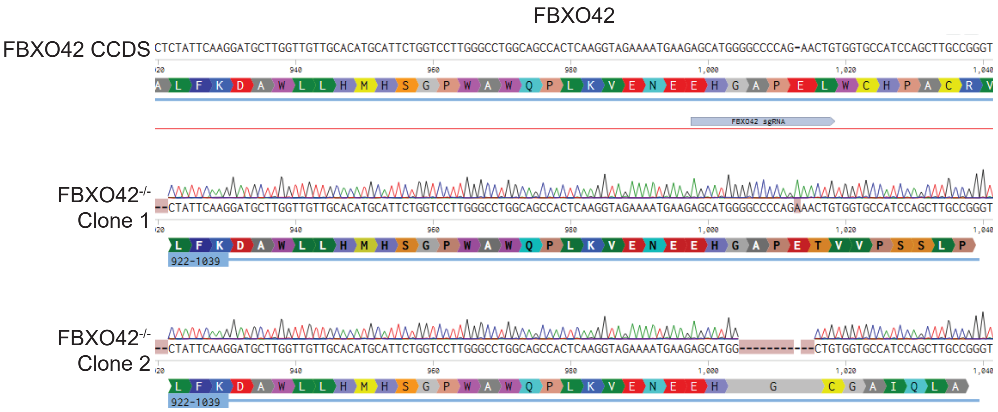

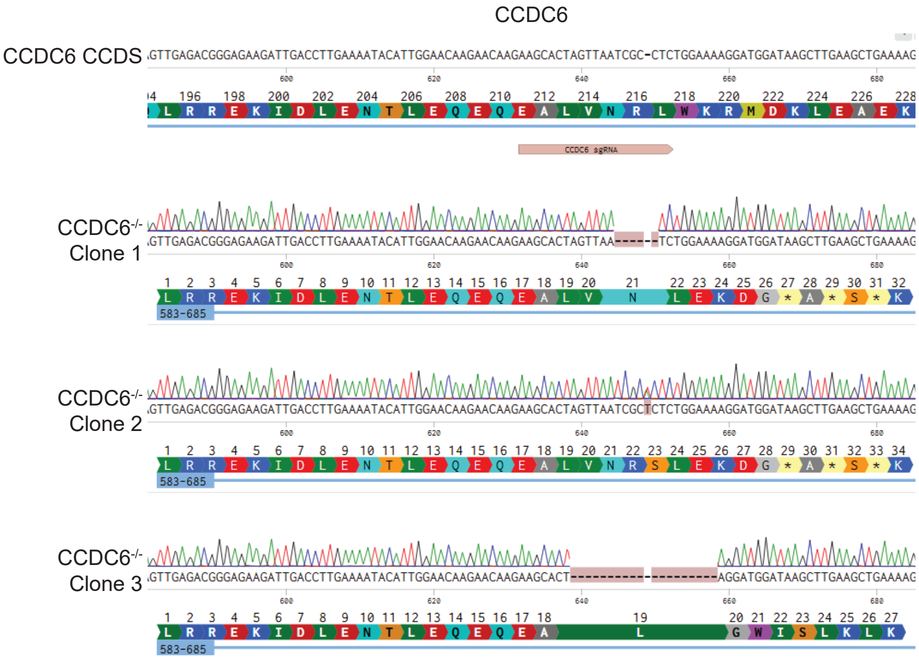

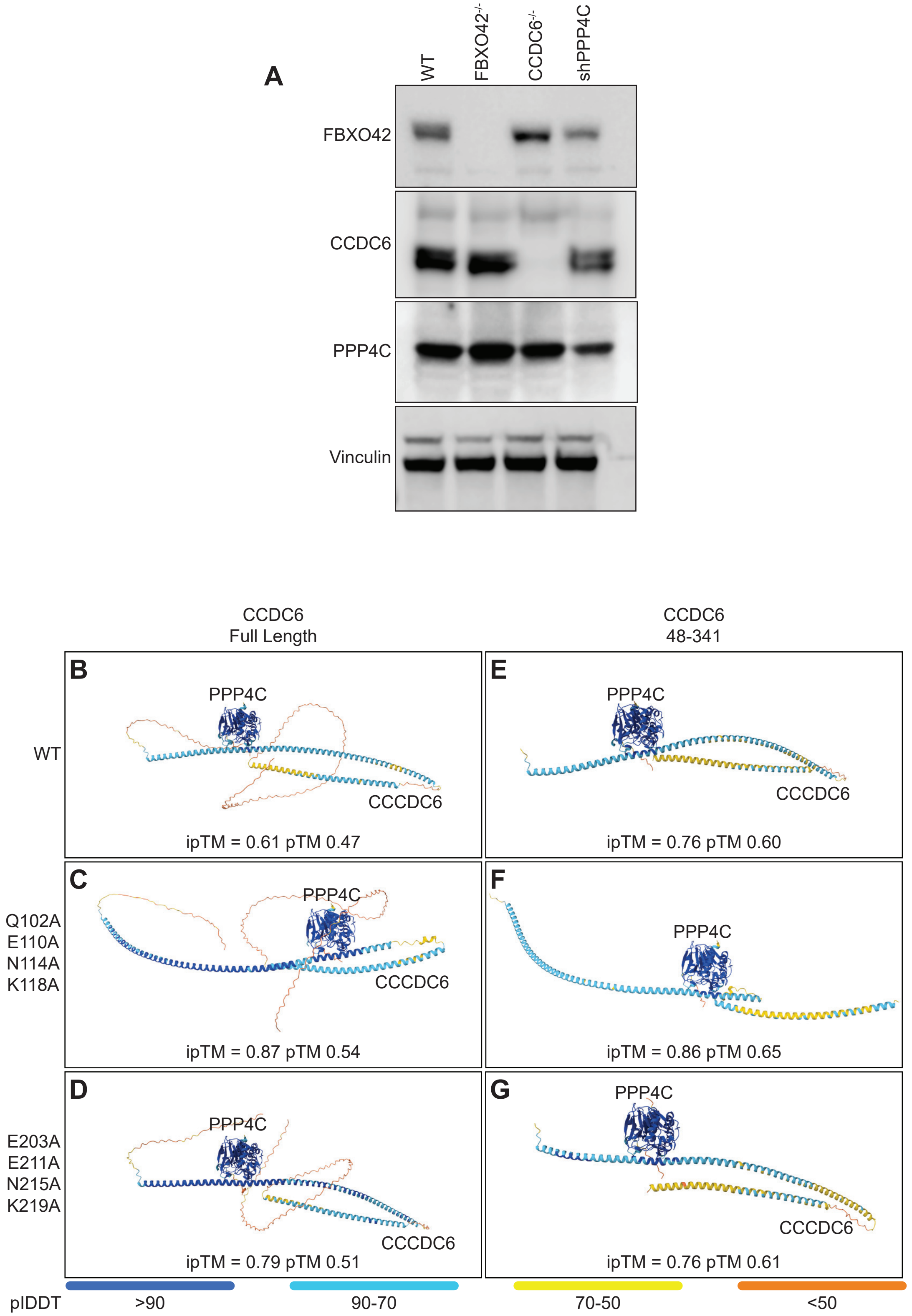

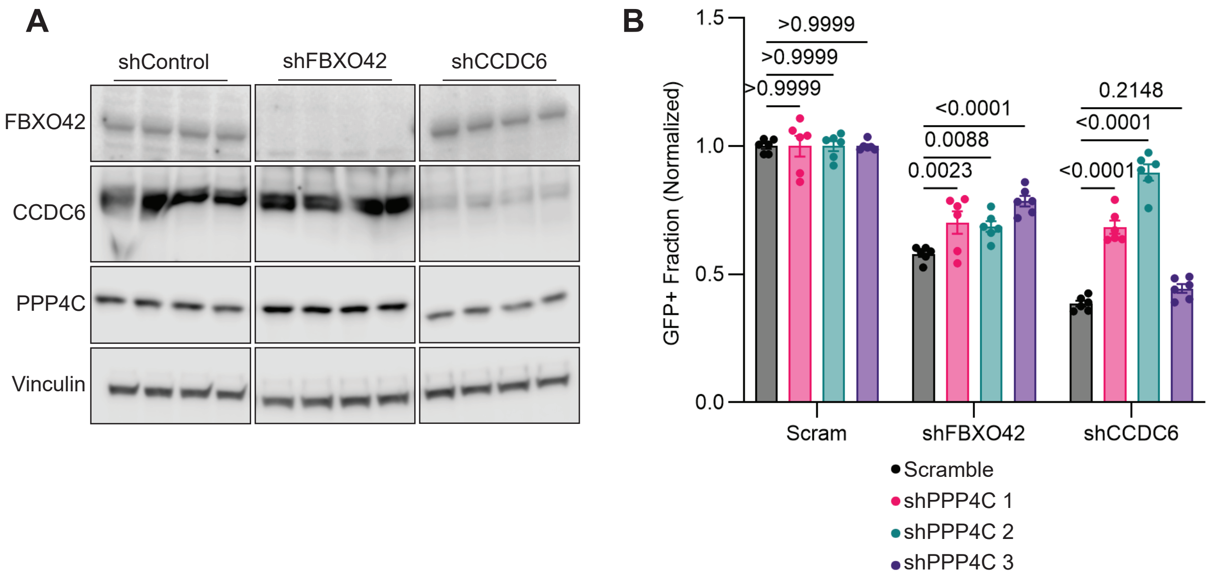
